# Pharmacogenomic diversity at tuberculosis-treatment loci across African populations: phased haplotypes, regulatory architecture and NAT2 acetylator inference

**DOI:** 10.64898/2026.11.22.768388

**Authors:** Ritah F. Mutagonda, Reuben S. Maghembe, Vesla I. Kullaya, Ananyo Choudhury, Collen Masimirembwa, Neil Martinson, Scott Hazelhurst, Michèle Ramsay, Jocelyn Gayenga, Scott Heysell, Stellah G. Mpagama

## Abstract

**Background:** Tuberculosis imposes a major public-health burden in Africa, while variation in isoniazid and rifampicin disposition creates opportunities for improved pharmacotherapy. The population architecture of extended, phased pharmacogenomic backgrounds remains less well characterized than individual marker frequencies.

**Methods:** We analyzed high-coverage, phased GRCh38 genomes from 661 unrelated individuals sampled from five continental-African and two African-diaspora populations at NAT2, SLCO1B1, CES2, HLA-B, NR1I2 and UGT1A1. We integrated frequency differentiation, reduced phased signatures, linkage disequilibrium, functional annotation and NAT2 core-allele inference with individual-clustered permutation tests.

**Results:** The six loci contributed 17,223 annotated variant records and 276 entries in a capped frequency-differentiation register. Exact NAT2 core-definition matching was achieved for 1,321/1,322 chromosomes (99.92%), and determinate genotype-inferred acetylator categories for 620/661 individuals (93.80%). Among the sampled continental-African populations, 35-site noncore background distributions differed within NAT2 *5 (BH-adjusted q=0.000100), *4 (q=0.024733) and *6 (q=0.000350) groups. Distinct phased and linkage architectures also characterized transporter, hydrolysis, regulatory and HLA-B genomic regions.

**Conclusion:** African population-associated variation extends beyond commonly used pharmacogenetic allele definitions. The observed NAT2 backgrounds and complementary locus signatures provide a foundation for targeted molecular studies and prospective pharmacokinetic validation that could inform future population-responsive TB treatment research.

## 1. Introduction

Tuberculosis (TB) remains a substantial global public-health challenge, with an estimated 10.7 million incident cases and 1.23 million TB-associated deaths in 2024; the WHO African Region accounts for a disproportionate burden (World Health Organization, 2025). Drug-susceptible TB is usually treated with isoniazid, rifampicin, pyrazinamide and ethambutol, yet interindividual differences in drug exposure, treatment-associated liver injury and comorbid conditions complicate therapy. Drug-induced toxicity may require treatment interruption, while inadequate exposure can compromise treatment efficacy and contribute to resistance selection; both motivate pharmacokinetic optimization studies. These concerns are especially relevant in African settings with high TB/HIV co-burden and heterogeneous clinical resources (McIlleron et al., 2015; Mouton et al., 2016; Abulfathi et al., 2019; Tostmann et al., 2008; Oelofse et al., 2023).

At the molecular level, isoniazid acetylation is influenced by arylamine N-acetyltransferase 2 (*NAT2*), whereas transport and hydrolysis processes involving *SLCO1B1* and *CES2* are candidate contributors to rifampicin disposition. *NAT2* genotype–phenotype associations and genotype-guided isoniazid dosing have been investigated clinically (Kinzig-Schippers et al., 2005; Azuma et al., 2013; Hein and Millner, 2021). African pharmacokinetic studies have also evaluated *SLCO1B1, CES2* and other host determinants of rifampicin exposure (Chigutsa et al., 2011; Dompreh et al., 2018; Naidoo et al., 2019). *NR1I2* encodes pregnane X receptor (PXR), an established regulator of xenobiotic-responsive transcription, including rifampicin-inducible pathways (Kliewer et al., 1998; Goodwin et al., 1999). *UGT1A1* promoter variation is an experimentally supported determinant of bilirubin glucuronidation (Bosma et al., 1995), while classical *HLA-B* variation offers a distinct immunogenetic route to investigating susceptibility to anti-TB drug-induced liver injury (Nicoletti et al., 2021).

A further challenge is the limited transferability of sparse marker panels across deeply structured African and admixed populations. African genomic diversity, local linkage disequilibrium and the spectrum of functional variants vary across sampled groups, as established by population-genomic and ADME-gene studies (Tishkoff et al., 2009; 1000 Genomes Project Consortium, 2015; da Rocha et al., 2021; Pereira et al., 2021). Broader representation in pharmacogenomic discovery and careful population-specific interpretation are priorities for equitable implementation (Chaudhry et al., 2016; Popejoy, 2019; Dandara et al., 2019). Published *NAT2* surveys demonstrate extensive geographic and haplotype diversity (Sabbagh et al., 2008; Mortensen et al., 2011), including variation in sub-Saharan African coding haplotypes (Patin et al., 2006). Nevertheless, most anti-TB pharmacogenetic comparisons focus on a limited set of alleles or individual variants; variation in the extended phased background after conditioning on the same *NAT2* core allele has been less explicitly investigated.

High-coverage phased genome resources now make it possible to resolve this biological question without relying exclusively on unphased single-marker summaries (Byrska-Bishop et al., 2022). We therefore investigated six predefined loci—NAT2, SLCO1B1, CES2, HLA-B, NR1I2 and UGT1A1—across five continental-African and two African-diaspora population samples. Specifically, we asked (i) how allele frequencies, reduced haplotype signatures and local LD differ between groups; (ii) which functional and regulatory features accompany this variation; how completely captured NAT2 definitions support acetylator inference; and (iii) whether common NAT2 core alleles occur on statistically different 35-site backgrounds. Connecting population haplotypes with allele-function annotations and a transparent translational evidence hierarchy can nominate precise candidates for future enzyme assays, hepatocyte studies and clinical pharmacokinetic research, without assuming that population-frequency differences demonstrate a causal treatment effect (Masiphephethu et al., 2022; Oelofse et al., 2023; Papanikolaou et al., 2026).

## 2. Materials and Methods

### 2.1 Study design and population sampling

We conducted a secondary analysis of publicly available 30-fold-coverage phased whole-genome sequencing data on GRCh38 from the 1000 Genomes Project (Byrska-Bishop et al., 2022). The analysis included 661 unrelated individuals: Esan in Nigeria (ESN, n=99), Gambian in Western Divisions in The Gambia (GWD, n=113), Luhya in Webuye, Kenya (LWK, n=99), Mende in Sierra Leone (MSL, n=85), and Yoruba in Ibadan, Nigeria (YRI, n=108), totaling 504 continental-African individuals. The comparative diaspora/admixture group comprised African Caribbeans in Barbados (ACB, n=96) and Americans of African Ancestry in SW USA (ASW, n=61), totaling 157 individuals. These labels refer to sampled population cohorts, rather than all individuals from their corresponding geographic communities. No new participant recruitment or specimen collection was performed.

### 2.2 Genomic regions and discovery scope

The wider discovery screen examined *NAT2, SLCO1B1, ABCB1, CES2, UGT1A1, NR1I2, HLA-B* and *NR1I3*. Locus coordinates were harmonized to GRCh38 and evaluated within gene-body and flanking intervals of 5, 25 and 100 kb. The principal population comparisons used a 25-kb extension on either side of each gene. This article reports six prespecified loci (*NAT2, SLCO1B1, CES2, HLA-B, NR1I2* and *UGT1A1*). Gene-level annotation counts reported for these loci were calculated from their respective records; summary counts from the eight-gene screen are distinguished explicitly.

### 2.3 Phased variant extraction and quality control

Phased GRCh38 chromosome VCFs were queried within each predefined locus. The original phased multiallelic representation was retained for haplotype analyses, while a split-allele representation supported variant-effect annotation. The broader eight-gene extraction workflow contained 16 source-by-region combinations and 48,744 source-preserved record observations across extraction rows. Sample identities, interval definitions, chromosome sources and checksums were tracked in manifests. BCFtools/SAMtools provides the relevant indexed VCF-processing framework (Danecek et al., 2021); however, the exact version and command-line history of each frozen extraction must be reconciled against the original execution logs rather than reconstructed from this manuscript.

### 2.4 Population-genomic analyses

Within each sampled population and locus, analyses summarized callability, allele frequency, heterozygosity, nucleotide diversity and phased completeness. Population differentiation used sites with call rate ≥0.95. Pairwise Hudson FST followed the ratio of summed sitewise numerators and denominators, retaining negative estimates as calculated rather than truncating them (Bhatia et al., 2013). A descriptive, capped frequency-differentiation register required a continental allele-frequency range ≥0.10 and retained no more than 50 entries per gene. Reduced common-SNP haplotype matrices required call rate ≥0.98 and minor-allele frequency ≥0.05; ambiguous multiallelic split positions were excluded. Haplotype- and LD-based population contrasts were interpreted in the context of documented African linkage and genetic structure (Tishkoff et al., 2009; Pereira et al., 2021).

### 2.5 Functional annotation and molecular-QTL triangulation

Variants were functionally annotated using the Ensembl Variant Effect Predictor (VEP), retaining available transcript, HGVS, MANE, coding, regulatory, predicted splicing and protein-feature fields (McLaren et al., 2016). MANE transcript designations provide a standardized reference between RefSeq and Ensembl annotations (Morales et al., 2022). Available information was arranged into candidate regulatory, RNA-processing and protein-related mechanisms. Selected signals were compared with GTEx Adult Analysis V10 eQTL and sQTL results as tissue-specific external context; GTEx Consortium (2020) provides foundational atlas evidence, whereas Adult Analysis V10 is the actual data-release provenance (GTEx Portal, Adult Analysis V10). Neither resource measures expression in the 661 individuals analyzed here.

### 2.6 Phased reduced signatures and population comparisons

Reduced phased signatures combined selected pharmacogenetic anchor sites, continental frequency-contrasting variants and annotated functional candidates. The source algorithm initially required ≥750 bp separation and at most 40 sites per gene. An abundance-supported signature required ≥5 continental chromosome observations and ≥2% frequency in at least one continental population. Similar signatures were clustered at a Hamming-distance threshold of the lesser of two differences or 5% of panel length. Signature diversity was summarized using Shannon effective numbers; differences between full signature-frequency distributions were quantified using Jensen–Shannon divergence (JSD), an information-theoretic measure defined for distributions with different support (Lin, 1991). Pairwise linkage disequilibrium was summarized as r^2^ within fixed ordered SNP panels.

### 2.7 *NAT2* alleles, diplotypes and phenotype inference

*NAT2* haplotypes were reconstructed from phased chromosome copies and matched to a frozen PharmVar/CPIC-linked capture of core allele definitions and functional assignments. Current *NAT2* nomenclature and functional uncertainty were interpreted against the curated gene-focused literature (Papanikolaou et al., 2026), earlier allele–phenotype evidence (Hein and Millner, 2021) and substrate-specific functional observations (Fukunaga et al., 2021). Three distinct endpoints were tabulated: (i) exact compatibility with a captured core definition, (ii) interpretability of the allele-function assignment and (iii) interpretability of the individual diplotype-derived acetylator category. Thirty-five sites outside the captured core definitions formed the phased noncore background. One unmatched chromosome was left without a newly invented star-allele designation.

Five common *NAT2* core groups (*5, *4, *6, *14 and *1) met the predefined continental-population representation criteria for conditional-background analysis. For each core group, generalized JSD compared the distributions of 35-site noncore backgrounds across five populations (Lin, 1991). Statistical significance was evaluated using 50,000 label permutations with carrier-level clustering; chromosomes belonging to the same individual were kept together. The Benjamini–Hochberg procedure controlled false discovery rate across five global tests, and a separate family of 50 pairwise tests was adjusted independently (Benjamini and Hochberg, 1995). This design tests population-associated distributional heterogeneity conditional on core allele, not differences in drug concentration.

### 2.8 *HLA-B* locus and clinical-evidence boundaries

*HLA-B* regional phased SNP and linkage patterns were analyzed at the sequence-marker level, not converted to classical *HLA-B* genotypes. Dedicated HLA allele assignment would require a separately validated high-resolution typing workflow and accepted allele nomenclature such as IPD-IMGT/HLA (Robinson et al., 2020). Clinical anti-TB HLA associations were examined as external literature rather than as outcomes observed in this genomic dataset (Nicoletti et al., 2021). The literature-evidence assessment was kept analytically separate from source variant tables.

### 2.9 Governance and reproducibility

The study retained public-data source identities and versioned local analysis and figure manifests. Reported analysis logic includes region-indexed VCF processing, VEP annotations, core-definition matching, r^2^ LD, Hamming-distance clustering, Shannon diversity, generalized JSD, individual-clustered permutation tests and Benjamini–Hochberg correction (Danecek et al., 2021; McLaren et al., 2016; Lin, 1991; Benjamini and Hochberg, 1995). The current source archive does not identify all historical Python/R package versions, exact VEP invocation settings or complete source-input checksums. These unresolved details require reconciliation against the frozen workflow records before final code release; no unverified version information has been added.

## 3. Results

### 3.1 Population panel and locus discovery framework

The analytical cohort comprised 661 unrelated individuals, including 504 individuals from five sampled continental-African populations and 157 from two diaspora/admixture populations (Table 1). All principal figures and locus-level findings refer to the prespecified six-gene manuscript subset unless an eight-gene discovery-stage result is explicitly identified. This separation establishes the denominators for population comparisons, variant summaries and phenotype-inference completeness.

**Table 1.**
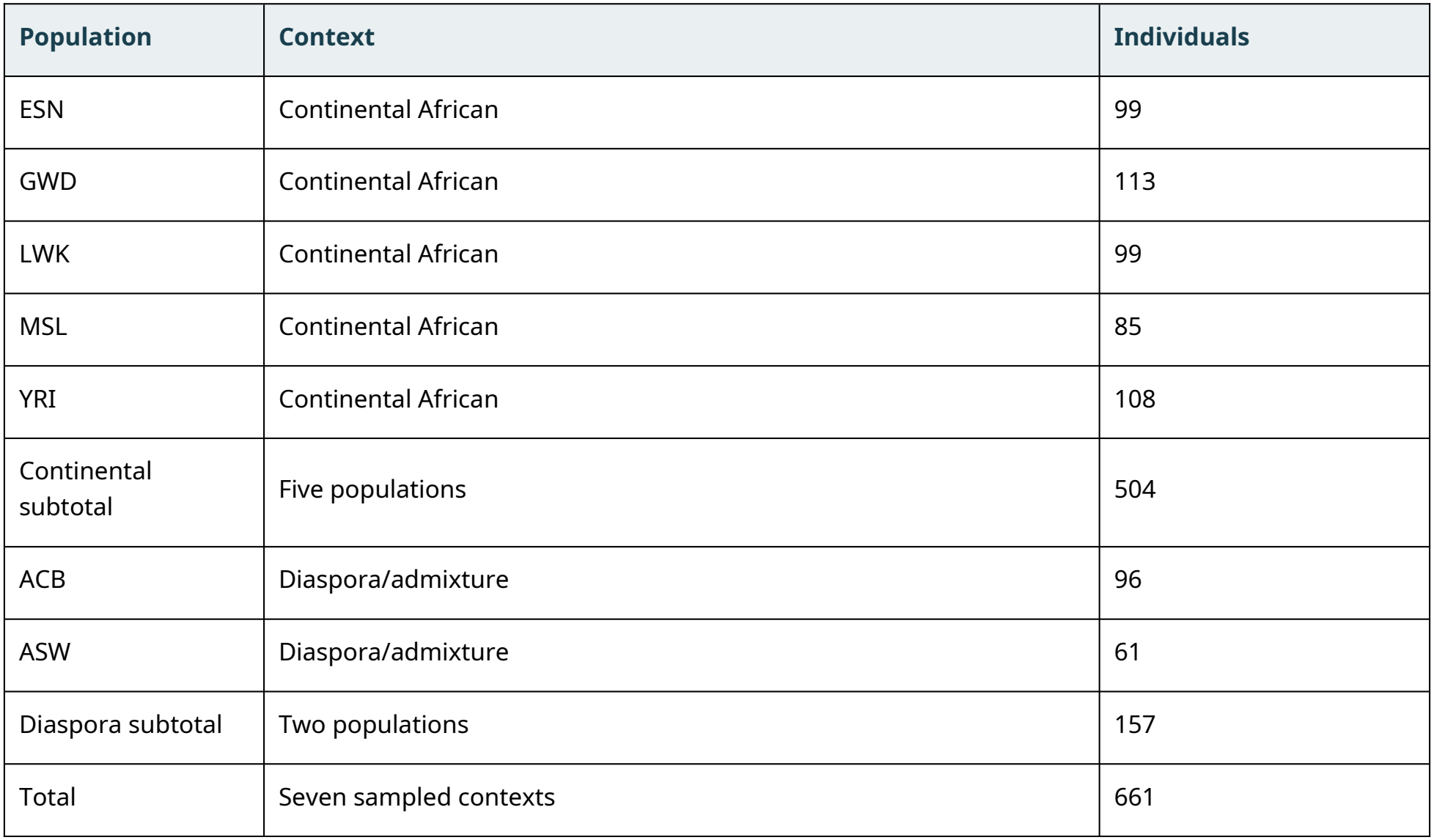
Populations and samples included in the genomic analysis. Population abbreviations: ESN, Esan in Nigeria; GWD, Gambian in Western Divisions, The Gambia; LWK, Luhya in Webuye, Kenya; MSL, Mende in Sierra Leone; YRI, Yoruba in Ibadan, Nigeria; ACB, African Caribbeans in Barbados; ASW, Americans of African Ancestry in the southwestern USA. Sample counts reflect the frozen unrelated-individual panel, not a clinical recruitment cohort; original source: Byrska-Bishop et al. (2022). Individual-level sex composition, HIV status and sampling dates are not included because they have not been authenticated for this analytic subset.

| **Population** | **Context** | **Individuals** |
|---|---|---|
| ESN | Continental African | 99 |
| GWD | Continental African | 113 |
| LWK | Continental African | 99 |
| MSL | Continental African | 85 |
| YRI | Continental African | 108 |
| Continental subtotal | Five populations | 504 |
| ACB | Diaspora/admixture | 96 |
| ASW | Diaspora/admixture | 61 |
| Diaspora subtotal | Two populations | 157 |
| Total | Seven sampled contexts | 661 |

### 3.2 Genetic variation and functional annotation

Across the six target loci, 17,223 variant records were annotated (Table 2). *SLCO1B1* contributed the largest annotation count (5,972), followed by *HLA-B* (4,104); the remaining counts were 2,089 at *NR1I2*, 1,963 at *NAT2*, 1,789 at *UGT1A1* and 1,306 at *CES2*. The capped prioritized register contained 276 entries meeting the allele-frequency range criterion; *HLA-B* showed the largest maximum single-variant population frequency range (0.3131). The wider eight-gene screen had 24,372 annotations and 376 prioritized entries and is not included in these six-gene totals.

**Table 2.** Variant annotations and prioritized differentiated-variant entries at the six target loci.

| **Locus** | **Annotated variants** | **Differentiated entries\*** | **Max AF range** |
|---|---|---|---|
| NAT2 | 1,963 | 50 | 0.199 |
| SLCO1B1 | 5,972 | 50 | 0.259 |
| CES2 | 1,306 | 27 | 0.234 |
| HLA-B | 4,104 | 50 | 0.313 |
| NR1I2 | 2,089 | 50 | 0.217 |
| UGT1A1 | 1,789 | 49 | 0.247 |
| Total | 17,223 | 276 | - |

\*Gene-level differentiated-variant entries: continental allele-frequency range ≥0.10, with a maximum of 50 retained per gene. The total is a capped selected register, not the number of all genuinely differentiated variants.

### 3.3 Phased signature architecture

The parent eight-gene discovery workflow yielded 309 reduced-signature sites, 10,576 chromosome-by-locus observations, 4,167 population-by-signature frequency entries, 1,798 differentiation entries, 194 major signatures and 126 clusters. These parent totals were not redefined as six-gene outputs. The manuscript-specific six-locus comparison instead highlighted ten contrasting phased-signature cluster families (Figure 1).

**Figure 1.**
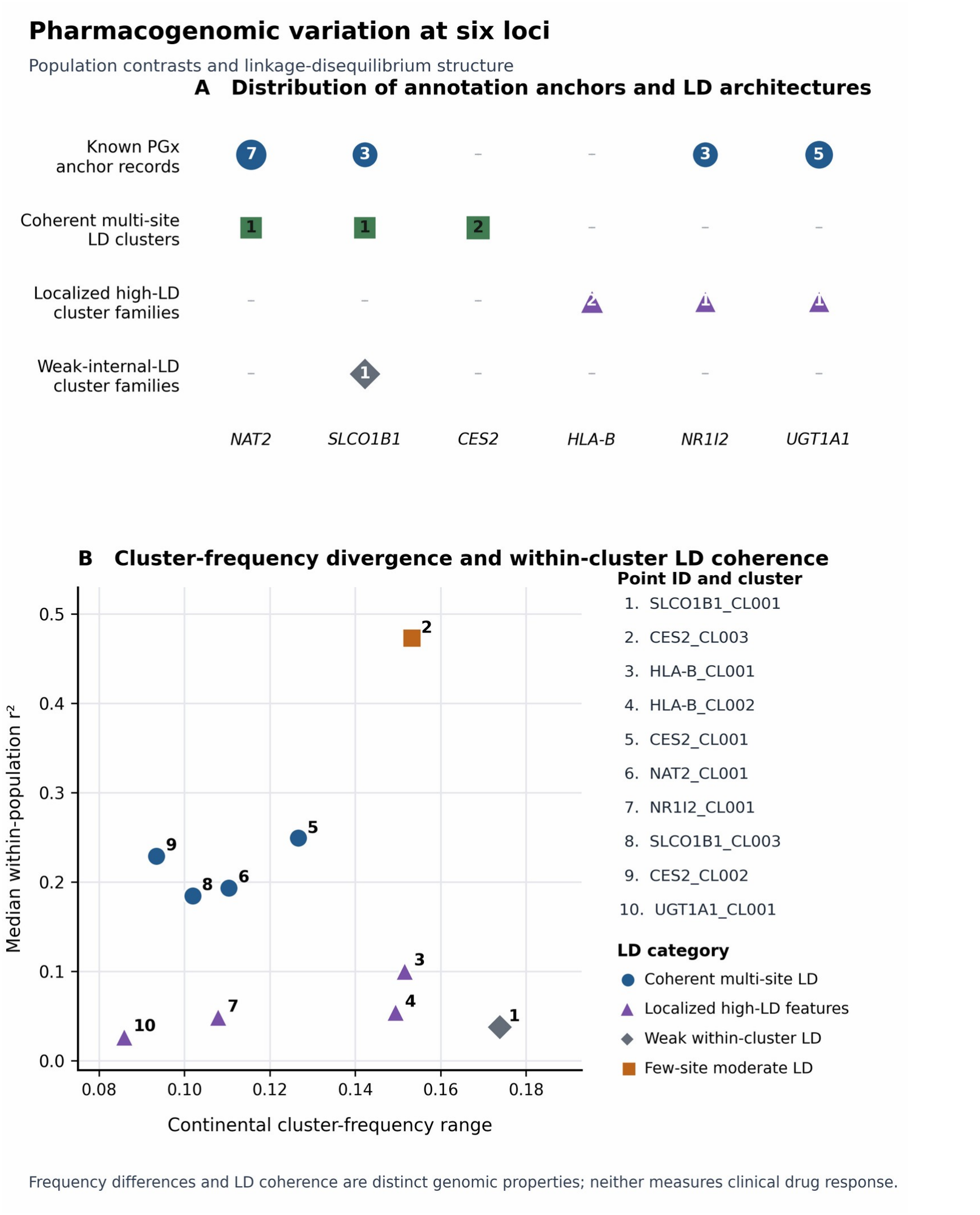
Phased genomic variation and population contrasts at six pharmacogenomic loci. (A) Curated pharmacogenomic-anchor counts and regional linkage architecture across six loci. (B) Continental population-frequency ranges and population-median within-cluster r^2^ for ten phased-signature cluster families. These quantities describe genomic variation, not treatment outcomes or classical HLA-B genotype assignments. Abbreviations: r^2^, squared correlation coefficient (linkage disequilibrium); LD, linkage disequilibrium; HLA, human leukocyte antigen; SNP, single-nucleotide polymorphism; NAT2, N-acetyltransferase 2; SLCO1B1, solute carrier organic anion transporter family member 1B1; CES2, carboxylesterase 2; NR1I2, nuclear receptor subfamily 1 group I member 2; UGT1A1, UDP glucuronosyltransferase family 1 member A1.

For the principal three-locus comparison, 40 *NAT2* sites, 40 *SLCO1B1* sites and 29 *CES2* sites were displayed in matched population-specific LD matrices across ESN, GWD, LWK, MSL and YRI (Figure 2). Identical ordered marker positions within each locus revealed regionally variable patterns of stronger and weaker pairwise marker correlation, generating candidates for more focused haplotype-aware comparison.

**Figure 2.**
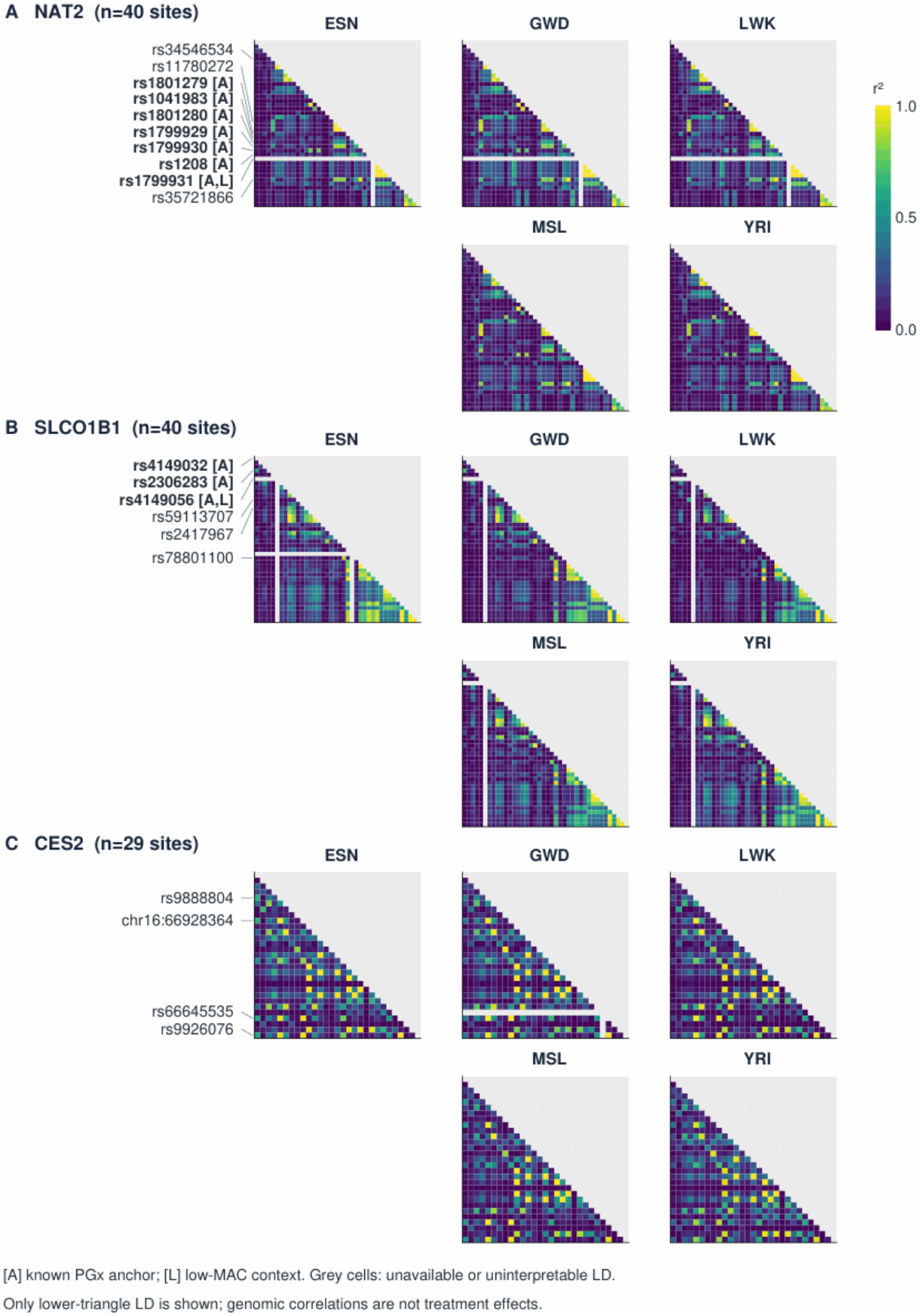
Population-specific linkage disequilibrium at *NAT2, SLCO1B1* and *CES2*. (A-C) Pairwise linkage disequilibrium (r-squared) at identical ordered SNP positions within NAT2 (40 sites), SLCO1B1 (40 sites) and CES2 (29 sites), respectively, among ESN, GWD, LWK, MSL and YRI. The five population matrices for each locus are arranged across two rows for legibility; each heatmap displays the lower triangle. Colour represents pairwise r-squared from 0 to 1; grey cells are unavailable or non-interpretable. [A] denotes a recognized pharmacogenomic anchor and [L] a low-minor-allele-count context. These are genomic, not treatment-outcome, comparisons. Abbreviations: ESN, Esan in Nigeria; GWD, Gambian in Western Divisions, The Gambia; LWK, Luhya in Webuye, Kenya; MSL, Mende in Sierra Leone; YRI, Yoruba in Ibadan, Nigeria; SNP, single-nucleotide polymorphism; LD, linkage disequilibrium; MAF, minor allele frequency.

### 3.4 Locus-prioritized population contrasts

Of ten highlighted six-locus cluster families, *SLCO1B1*_CL001 exhibited the largest continental between-population frequency range (0.1737), reaching a within-MSL frequency of 0.3000 (Figure 1). These differences illustrate why a multi-site signature enriched in one sampled population may provide a more informative experimental construct than its individual variants considered in isolation.

Figure 3 extends this comparison to three mechanistically distinct regions: *HLA-B* regional SNP-LD islands, 60 *NR1I2* anchor-pair/population LD values and 25 *UGT1A1* anchor-profile records centered on rs3064744. Together, these displays identify candidate immunogenetic-region and regulatory haplotype contexts without assigning a clinical effect to the local marker correlations.

**Figure 3.**
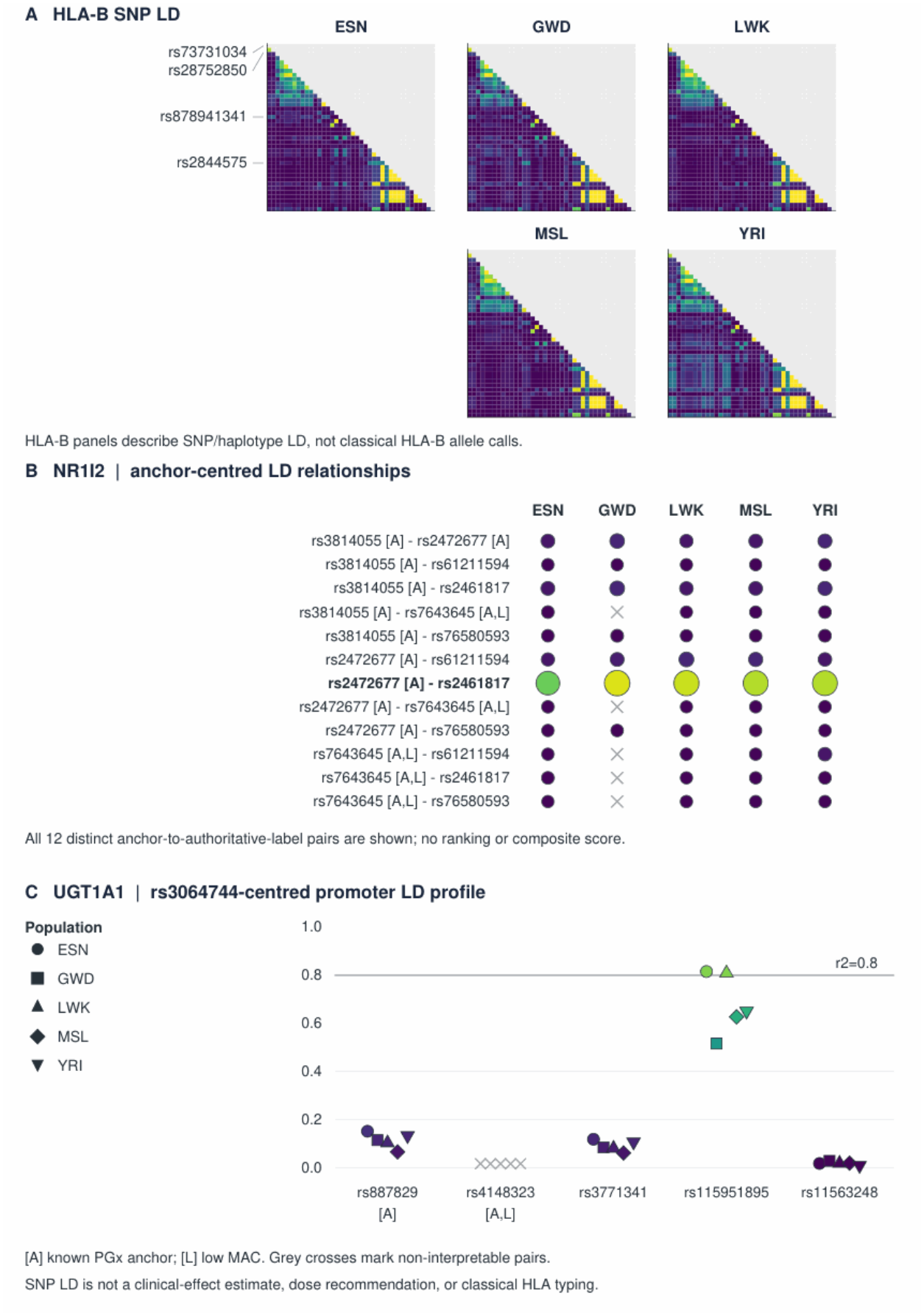
Local linkage architecture at *HLA-B, NR1I2* and *UGT1A1*. (A) Population-specific SNP linkage disequilibrium in the HLA-B region across the five continental-African populations; this is not classical HLA-B typing. (B) Twelve distinct NR1I2 anchor-to-marker pairs, with colour and marker area encoding pairwise r-squared; grey crosses indicate non-interpretable values. (C) Population-specific pairwise r-squared between UGT1A1 rs3064744 and five promoter-region partners. The horizontal reference corresponds to r-squared = 0.8. These panels characterize sequence-variation architecture, not clinical effects. Abbreviations: HLA-B, human leukocyte antigen B; NR1I2, nuclear receptor subfamily 1 group I member 2; UGT1A1, UDP glucuronosyltransferase family 1 member A1; SNP, single-nucleotide polymorphism; LD, linkage disequilibrium; r^2^, squared correlation coefficient.

### 3.5 *NAT2* phased alleles and inferred acetylator phenotypes

Across all seven sampled populations, 1,321 of 1,322 phased *NAT2* chromosomes (99.92%) matched a captured core definition exactly. Increased- or decreased-function status was resolved for 1,280 chromosomes (96.82%); 41 chromosomes had uncertain function and one was unmatched. Consequently, 620 of 661 individuals (93.80%) received determinate genotype-inferred acetylator categories, while 41 remained indeterminate (Table 3).

**Table 3.** NAT2 captured-definition matching and phenotype interpretability by sampled population. Table 3 is retained separately from Table 1 because it reports NAT2 inference endpoints and completeness rather than recruitment or population-panel composition.

| **Population** | **Individuals** | **Exact core calls** | **Determinate** | **Determinate %** | **Indeterminate** |
|---|---|---|---|---|---|
| ESN | 99 | 198/198 | 93 | 93.9% | 6 |
| GWD | 113 | 225/226 | 98 | 86.7% | 15 |
| LWK | 99 | 198/198 | 94 | 94.9% | 5 |
| MSL | 85 | 170/170 | 83 | 97.6% | 2 |
| YRI | 108 | 216/216 | 101 | 93.5% | 7 |
| ACB | 96 | 192/192 | 91 | 94.8% | 5 |
| ASW | 61 | 122/122 | 60 | 98.4% | 1 |
| Total | 661 | 1321/1322 | 620 | 93.8% | 41 |

Among continental-African participants, exact matches covered 1,007/1,008 chromosomes (99.90%), while 469/504 people (93.06%) received determinate inferred acetylator phenotypes. The highest proportion of indeterminate individuals occurred in GWD (15/113). The sole unmatched phased core combination was HG03025 haplotype 2 in GWD, associated with noncore background *NAT2*_BG35_c9366a061b58; an unmatched captured definition does not itself establish a novel functionally characterized allele.

Conditioning on core *NAT2* categories revealed considerable diversity of 35-site backgrounds in the 504-person continental panel (Table 4; Figure 4). The *5 group comprised 285 chromosomes on 34 noncore backgrounds, *4 comprised 233 on 44, and *6 comprised 182 on 27. Global population heterogeneity was statistically supported for *5 (generalized JSD=0.4657; BH q=0.000100), *4 (JSD=0.5519; q=0.024733) and *6 (JSD=0.5977; q=0.000350); *14 and *1 did not meet the adjusted significance threshold. Four adjusted pairwise contrasts for *5 and two for *6 were significant, while no individual *4 contrast survived the separate 50-test family. The overall and pairwise tests evaluate different hypotheses under different multiplicity corrections.

**Table 4.**
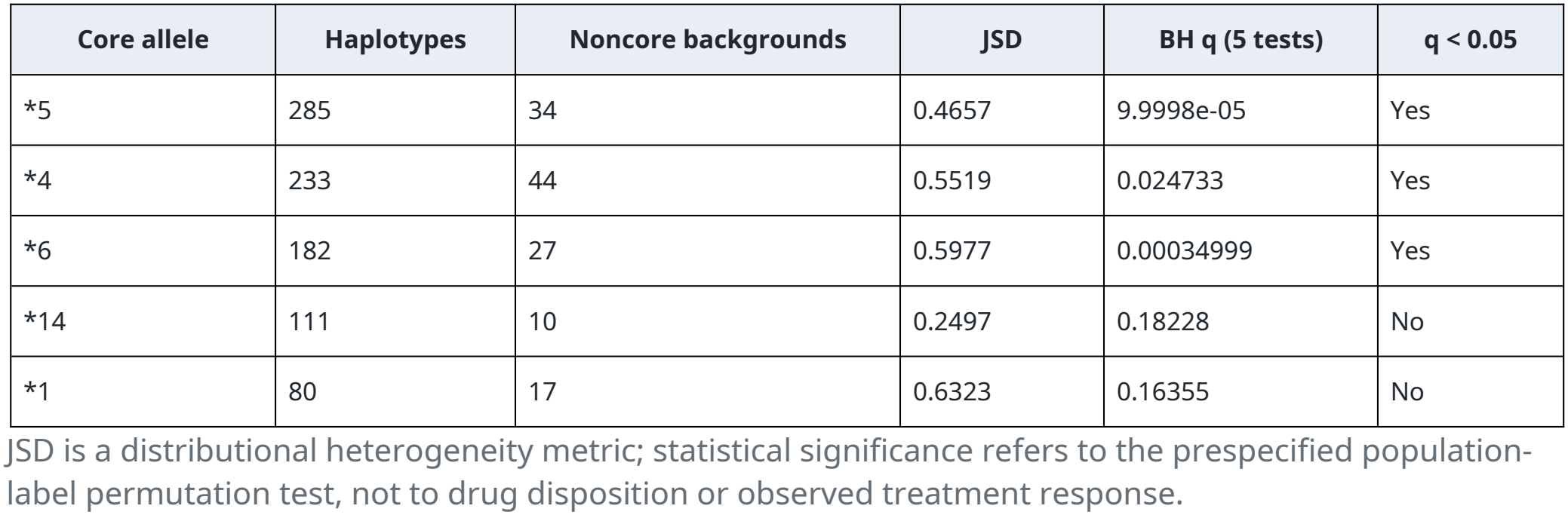
*NAT2* core-conditional background heterogeneity among continental-African populations.

| Core allele | Haplotypes | Noncore backgrounds | JSD | BH q (5 tests) | q < 0.05 |
|---|---|---|---|---|---|
| *5 | 285 | 34 | 0.4657 | 9.9998e-05 | Yes |
| *4 | 233 | 44 | 0.5519 | 0.024733 | Yes |
| *6 | 182 | 27 | 0.5977 | 0.00034999 | Yes |
| *14 | 111 | 10 | 0.2497 | 0.18228 | No |
| *1 | 80 | 17 | 0.6323 | 0.16355 | No |

JSD is a distributional heterogeneity metric; statistical significance refers to the prespecified population-label permutation test, not to drug disposition or observed treatment response.

**Figure 4.**
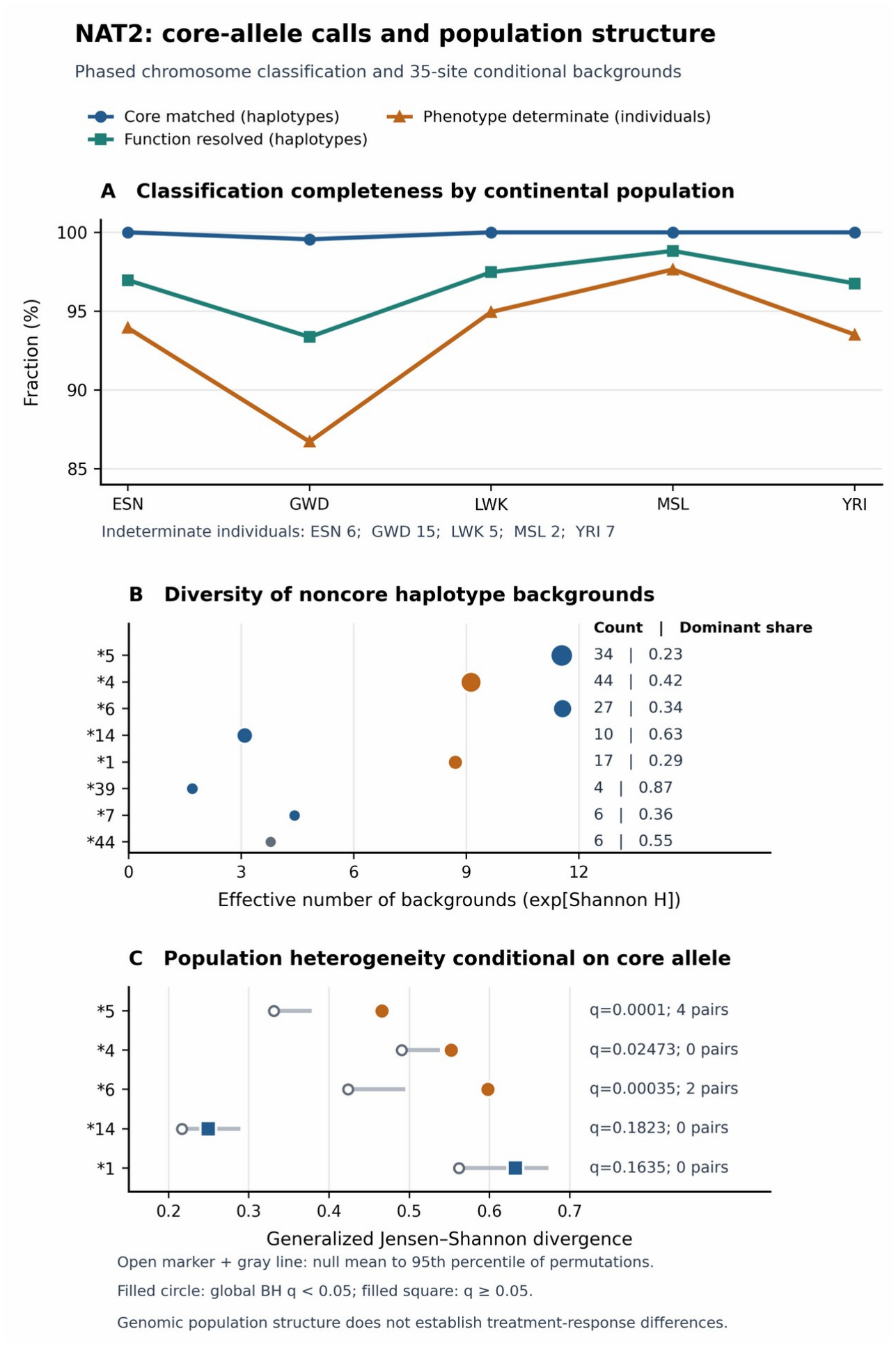
*NAT2* genotype-definition completeness and conditional haplotype backgrounds. (A) Exact NAT2 core-definition calls, functional assignment and determinate inferred acetylator phenotypes in the five continental-African populations. The expanded y-axis emphasizes interpretability differences; GWD contains the sole unmatched chromosome. (B) Shannon effective number of 35-site backgrounds for eight captured NAT2 core-allele groups; point sizes reflect continental chromosome counts and labels report distinct backgrounds and predominant background frequencies. (C) Generalized Jensen-Shannon divergence among population-specific noncore backgrounds conditional on allele group. Filled circles indicate global BH q < 0.05 across five tests; permutation-null means and percentile intervals are shown. Pairwise significance was controlled separately across 50 comparisons. The analysis concerns genomic population structure, not measured drug-response differences. Abbreviations: NAT2, N-acetyltransferase 2; GWD, Gambian in Western Divisions, The Gambia; BH, Benjamini–Hochberg; q, false-discovery-rate-adjusted p value; JSD, Jensen–Shannon divergence.

### 3.6 Regulatory triangulation and *HLA-B* interpretation

The parent eight-gene GTEx Adult V10 screen yielded 8,762 target-gene eQTL/sQTL association records, with no separately established six-locus record total. These external regulatory annotations offer candidate tissue-context links for molecular follow-up (GTEx Consortium, 2020). Within the *HLA-B* region, local LD architectures and reduced signatures including *HLA-B*_CL001/CL002 differed among sampled continental populations, motivating classical allele typing and immune-mechanism research as subsequent rather than completed steps.

### 3.7 Translational evidence assessment

Independent pharmacogenetic evidence identifies feasible validation endpoints for the observed genomic candidates. African studies have linked *NAT2* genetic variation with isoniazid disposition (Mthiyane et al., 2020; Masiphephethu et al., 2022), tested rifampicin pharmacokinetic correlates of *SLCO1B1*/*CES2* variation (Chigutsa et al., 2011; Dompreh et al., 2018; Weiner et al., 2010; Mukonzo et al., 2020), and investigated classical HLA associations with anti-TB drug-induced liver injury (Nicoletti et al., 2021). Those patient-level studies anchor experimentally testable questions without being treated as direct clinical validation of the current 661-genome signatures.

## 4. Discussion

This study identifies a previously underappreciated layer of African anti-tuberculosis pharmacogenomics: near-complete core *NAT2* allele matching coexists with extensive, statistically population-associated variation in the surrounding phased backgrounds. Across six loci, 17,223 annotated variant records, 276 selected differentiated entries and distinct population-specific linkage patterns describe a richer substrate for pharmacological hypothesis generation than isolated sentinel SNPs. Particularly striking is the heterogeneity of *NAT2* *5, *4 and *6 backgrounds even after conditioning on the same core allele group. This finding may imply that individuals sharing an acetylator allele category can nevertheless differ in linked local regulatory or structural features. The observation provides a specific mechanistic question—whether extended background modifies gene expression or enzymatic behaviour within the same core genotype—that can now be tested experimentally.

NAT2 offers the most direct translational bridge because it is involved in isoniazid acetylation, for which functional genetic variation affects exposure and has motivated genotype-informed dosing research (Kinzig-Schippers et al., 2005; Azuma et al., 2013; Hein and Millner, 2021). Our 99.92% exact core-definition matching indicates strong capture of recognized allele-defining patterns in this dataset, providing a useful starting point for pharmacogenomic panel development. Nevertheless, 41/661 individuals (6.20%) lacked a determinate inferred phenotype and the *5, *4 and *6 core groups contained 34, 44 and 27 distinct noncore backgrounds, respectively. Evidence from Ethiopian genotype–environment interaction and Tanzanian and South African research connects NAT2 variation to NAT2-dependent acetylation phenotypes relevant to isoniazid biotransformation, exposure and clearance, while experimental work demonstrates substrate-specific allele-function differences (Aklillu et al., 2018; Mthiyane et al., 2020; Masiphephethu et al., 2022; Fukunaga et al., 2021). Our observation therefore raises the possibility that subgroups, defined by an extended phased background, may differ in NAT2 expression or residual enzyme function even when their core call is identical. This hypothesis would be best tested by comparing matched core-allele carriers across backgrounds in hepatocyte expression assays, recombinant enzyme acetylation experiments and repeated-dose isoniazid pharmacokinetic studies. Such analyses could identify whether extended haplotypes add predictive information beyond established core alleles, which would be valuable for the design of parsimonious but population-appropriate genotyping assays (Verma et al., 2021; Rens et al., 2020).

Population differences in the transport and hydrolysis genes offer a second route to pharmacokinetic prioritization. *SLCO1B1* accounted for 5,972 annotated variants, and the *SLCO1B1*_CL001 signature reached a frequency of 0.3000 in the MSL sample while exhibiting a continental frequency range of 0.1737. Such a population-enriched signature may capture correlated marker combinations that are not adequately represented by a single widely typed *SLCO1B1* SNP. South African studies identified associations between selected *SLCO1B1* alleles and lower rifampicin concentrations, although replication and effect sizes vary across settings (Chigutsa et al., 2011; Weiner et al., 2010; Naidoo et al., 2019). A Ugandan pharmacogenetic study and a systematic review likewise identify both promise and heterogeneity in the literature on rifampicin exposure (Mukonzo et al., 2020; Sileshi et al., 2022). At *CES2*, the 1,306 annotated records and distinct LD profiles make it possible to nominate specific hydrolase-related backgrounds for comparison: experimental and clinical studies have investigated *CES2* variants alongside rifampicin disposition (Song et al., 2013; Dompreh et al., 2018; Thomas et al., 2020). We propose testing whether the observed haplotype configurations track with transporter uptake, hydrolase activity or rifampicin concentration–time profiles after accounting for dose, body size, HIV status and concomitant therapy (Abulfathi et al., 2019; Tostmann et al., 2013).

The regulatory loci *NR1I2* and *UGT1A1* demonstrate how phased genomic context can sharpen the design of mechanistic assays. *NR1I2* encodes PXR, a xenobiotic sensor implicated in drug-responsive transcription; rifampicin activation of the human PXR-linked CYP3A4 regulatory pathway has been demonstrated in reporter and expression systems (Kliewer et al., 1998; Goodwin et al., 1999). The observed *NR1I2* anchor-centred LD patterns could therefore guide the selection of regulatory haplotypes for PXR expression, promoter/enhancer reporter and transcriptomic assays. At *UGT1A1*, the rs3064744-centred promoter profiles are biologically interesting because promoter sequence can modify *UGT1A1* transcription and bilirubin conjugation (Bosma et al., 1995). The present profiles nominate linked promoter-region constructs worth comparing, rather than showing which exact marker is causal. Integration with GTEx tissue-dependent molecular-QTL signals and consistent MANE transcript annotation could refine this prioritization (GTEx Consortium, 2020; Morales et al., 2022). If a coherent phased background predicts altered reporter activity or transcript output under relevant drug exposure, that would supply an experimental basis for prospective study in hepatic models.

The distinct *HLA-B* regional signatures introduce an immunogenetic dimension to drug safety research. Figure 3 demonstrates localized differences in *HLA-B* region SNP correlation among the five continental-African samples, while published multi-ancestry research has associated a specific classical HLA allele with liver injury during isoniazid-containing TB treatment (Nicoletti et al., 2021). This convergence suggests a useful pathway: high-resolution *HLA-B* typing, concordance testing against phased regional markers, and immune-response studies could determine whether the observed population-enriched backgrounds tag biologically meaningful classical alleles. Because clinically interpreted HLA alleles are defined by a specialized polymorphic reference nomenclature, an HLA genotype should be assigned directly using a validated method, not inferred by naming local SNP clusters (Robinson et al., 2020). Linking the typed alleles to rigorously defined hepatic adverse-event phenotypes and immune mechanisms would translate the genomic maps into a focused, empirically assessable immunogenetic research programme.

A broader implication concerns pharmacogenomic portability and equitable study design. Deep population history, demographic structure, selection and recombination can influence African haplotype diversity (Tishkoff et al., 2009; Sabbagh et al., 2008; Mortensen et al., 2011). The present analysis shows why a core pharmacogenetic designation and the broader sequence context are complementary dimensions: the former supports interpretable standardization, whereas the latter exposes population-specific combinations to test. This is particularly important for admixed groups and for the development of panels expected to work across genetically diverse African settings (Pereira et al., 2021; da Rocha et al., 2021; Popejoy, 2019). Rather than implying that an entire named population shares a fixed drug-response trait, the data support selecting individuals and cell constructs by directly measured haplotypes. Such a strategy fits with African pharmacogenomics capacity-building and the proposed expansion of clinically relevant allele resources (Chaudhry et al., 2016; Dandara et al., 2019; Oelofse et al., 2023). The observed LD and frequency patterns are descriptive: distinguishing selection from demographic history and recombination would require dedicated, appropriately controlled evolutionary analyses not performed here.

The immediate translational opportunity is a prospective, genotype-stratified pharmacokinetic and functional-validation programme. Participants would require documented isoniazid dose, dosing schedule, dose-to-sample interval, adherence and repeated concentration–time sampling, together with NAT2 core diplotypes and extended phased backgrounds. In settings where relevant, analyses should prespecify antiretroviral regimens and potential drug–drug interactions, HIV status, and pregnancy versus postpartum state; pregnancy-associated physiological changes and postpartum recovery should not be treated as interchangeable with the nonpregnant state (Gausi et al., 2021; McIlleron et al., 2015; Malki and Pearson, 2020). Tanzania already has cohort-based evidence motivating NAT2 genotyping for individualized isoniazid dosing and pharmacokinetic investigation (Masiphephethu et al., 2022; Tostmann et al., 2013). The primary testable endpoint would be whether noncore NAT2 backgrounds explain additional variability in isoniazid clearance or exposure after accounting for core acetylator category and major covariates. Related but distinct mechanistic work could evaluate SLCO1B1-mediated uptake, CES2-mediated hydrolysis, PXR-dependent transcription and typed HLA alleles. The potential value of an expanded pharmacogenomic panel should be quantified prospectively through incremental predictive accuracy, discrimination or calibration, and ultimately clinically meaningful outcomes, rather than inferred from haplotype differences alone. These are proposed studies, not clinical or pharmacokinetic observations generated here.

### 4.1 Study limitations

The seven 1000 Genomes cohorts sample only a portion of continental-African and diaspora diversity, and these sample labels cannot be equated with population-wide clinical probabilities. The differentiated-variant register was capped at 50 entries per locus, and some parent summaries derive from an eight-gene rather than a six-gene screen. Functional annotations and GTEx molecular-QTLs were external and were not measured in these participants. No isoniazid or rifampicin concentrations, enzyme activity, clinical outcomes or liver-injury events were collected; classical HLA-B typing was not performed. Exact versions of some historical computational tools and a complete deposited reproduction environment remain to be recovered. These constraints define the appropriate next experimental and prospective steps. The source data were not a newly recruited clinical cohort; pregnancy, HIV status, treatment history and sex-stratified clinical metadata should not be asserted for individual samples without authenticated source metadata. Accordingly, study eligibility, demographic variables and sampling dates are limited to what can be documented from the source-project records.

## 5. Conclusions

In 661 publicly sourced phased genomes, six tuberculosis-treatment-relevant loci showed differentiated and complex local genomic architectures. Although *NAT2* core-definition assignments captured 99.92% of chromosomes, population-associated 35-site backgrounds persisted within common *NAT2* *5, *4 and *6 groups. These findings identify a new, directly testable question about how pharmacogenomic effects may vary across extended haplotypes within established allele categories. The complementary *SLCO1B1, CES2, NR1I2, UGT1A1* and *HLA-B* regional patterns further nominate transport, hydrolysis, transcriptional and immune-genetic targets for investigation. Functional assays and appropriately powered African TB pharmacokinetic cohorts can now evaluate which genomic backgrounds contribute measurable biological effects, strengthening the evidence base for future equitable genotype-informed treatment research.

## Data availability, ethics and declarations

### Data availability

Public, phased GRCh38 whole-genome resources are available through the International Genome Sample Resource 1000 Genomes 30x collection (https://www.internationalgenome.org/data-portal/data-collection/1000genomes_30x; Byrska-Bishop et al., 2022). Molecular-QTL comparisons refer to the GTEx Portal Adult GTEx Analysis V10 release (https://gtexportal.org/home/downloads/adult-gtex/; foundational atlas: GTEx Consortium, 2020). Derived locus and NAT2 summaries accompany this manuscript in Supplementary Tables S1–S6. A fully executable code/data deposit, including source-file checksums, captured core allele definitions and historical software environments, remains to be completed and will require an archival identifier before a public code-availability claim is made.

### Ethics statement

This work constitutes secondary analysis of publicly available, de-identified genomic resource data. No new participants were recruited and no new biospecimens or clinical outcomes were collected. Original participant consent and study governance are documented in the source resource publications (1000 Genomes Project Consortium, 2015; Byrska-Bishop et al., 2022).

## Author contributions

RFM: Conceptualization, Writing – original draft, Writing – review and editing. RSM: Conceptualization, Methodology, Data curation, Formal analysis, Software, Visualization, Writing – original draft, Writing – review and editing. VIK: Methodology, Writing – review and editing. AC, CM, NM, SHa (Scott Hazelhurst), MR and JG: Conceptualization, Resources, Writing – review and editing (provisional). SHe (Scott Heysell) and SGM: Conceptualization, Supervision, Project administration, Funding acquisition, Writing – review and editing (provisional). All manuscript-specific CRediT roles and final authorship approval require individual confirmation before submission.

## Acknowledgments

The authors acknowledge the participants and research teams who contributed to the 1000 Genomes and GTEx resources. The authors gratefully acknowledge the University of the Witwatersrand (Wits), Johannesburg, South Africa, for access to high-performance computing (HPC) resources that supported the computationally intensive genomic analyses undertaken in this study. ChatGPT (GPT-6 model; OpenAI, accessed October 2026) assisted with editorial organization, scientific-language revision, reference discovery and document assembly. No new genomic analyses were conducted with the language model. The authors are responsible for factual accuracy, verification of cited literature and approval of the final manuscript.

## Supplementary material

Supplementary Figure S1 shows inferred NAT2 acetylator-category frequencies. Machine-readable Supplementary Tables S1–S4 provide source locus annotations, inferred NAT2 phenotype frequencies, NAT2 background heterogeneity and population completeness; S5 provides the common-core integrated summary and S6 provides all 50 NAT2 pairwise population comparisons. Supplementary Table S1 retains the original eight-locus discovery-panel context, whereas the manuscript’s main tables report the six-locus subset.

