## Supplemental Data 1 for "Pharmacogenomic diversity at tuberculosis-treatment loci across African populations: phased haplotypes, regulatory architecture and NAT2 acetylator inference"

R7 preprint Version 1 | Consolidated source-preserving supplementary document | 22 November 2026

Contents: Supplementary Figure S1 and the complete underlying data of Supplementary Tables S1-S6. Table panels repeat identifier columns so all source fields remain readable and cross-referenceable.

These tables report genomic variant annotations, predicted acetylator categories and conditional background statistics. They do not represent measured drug concentrations, clinical outcomes or classical HLA allele typing.

| Component | Data source | Full data rows | Source columns |
|---|---|---|---|
| Figure S1 | Supplementary_Figure_S1.png | - | - |
| Table S1 | RITAH_Supplementary_Table_S1_Locus_Annotation_8GeneDiscovery.tsv | 8 | 11 |
| Table S2 | RITAH_Supplementary_Table_S2_NAT2_Acetylator_Frequencies.tsv | 28 | 6 |
| Table S3 | RITAH_Supplementary_Table_S3_NAT2_Background_Heterogeneity.tsv | 5 | 17 |
| Table S4 | RITAH_Supplementary_Table_S4_NAT2_Population_Completeness.tsv | 10 | 21 |
| Table S5 | RITAH_Supplementary_Table_S5_NAT2_Core_Integrated_Summary.tsv | 8 | 15 |
| Table S6 | RITAH_Supplementary_Table_S6_NAT2_Pairwise_50_Comparisons.tsv | 50 | 22 |

The original exact-precision data files are provided separately in the source preprint package. This PDF is a complete readable consolidation of their rows and columns, not a new analysis.

#### Supplementary Figure S1

Distribution of genotype-inferred NAT2 acetylator phenotypes across seven sampled population contexts. The bar-chart segments show rapid, intermediate, poor and indeterminate categories and do not represent observed clinical drug metabolism.

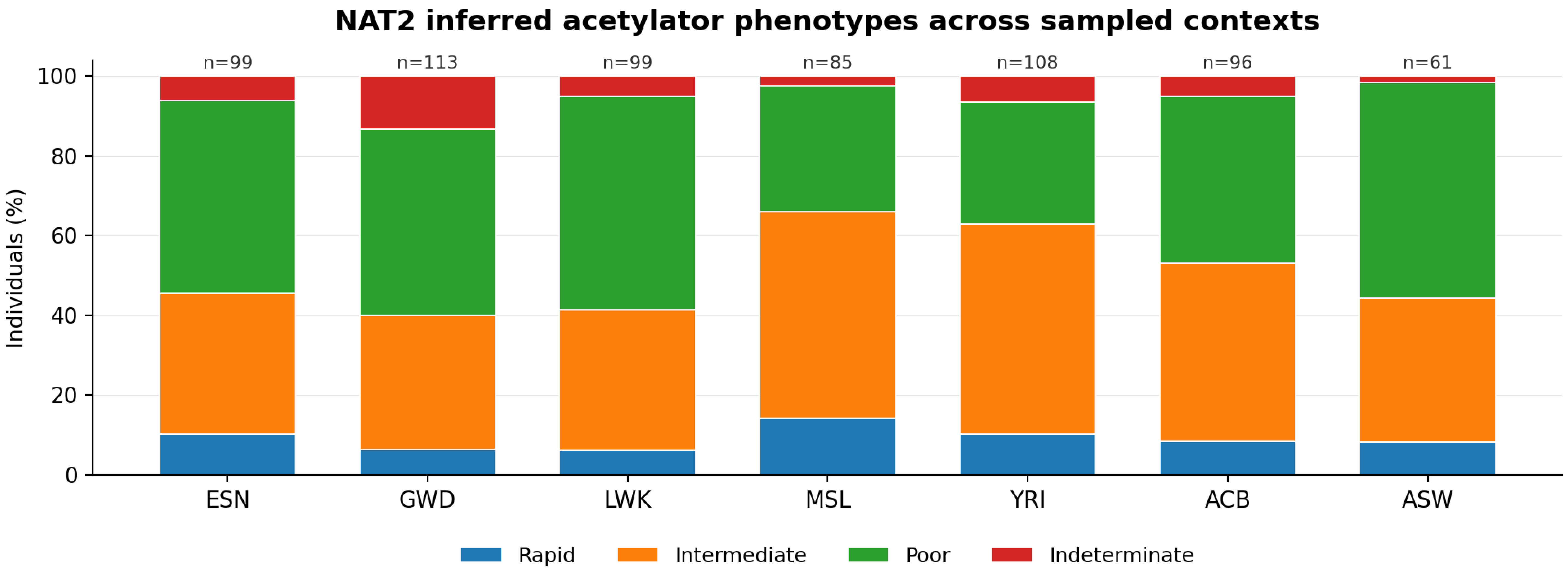


Population labels: ESN, GWD, LWK, MSL, YRI (continental African samples); ACB and ASW (African-diaspora samples). Values are genotype-inferred, not clinically measured.

### Supplementary Table S1. Eight-gene discovery-stage locus annotation summary

Source Table S1 is from the eight-gene parent discovery screen. Do not substitute these counts for the six-gene manuscript totals.

#### Supplementary Table S1 - panel A

| gene_symbol | all_unique_variants | differentiated_variants_AF_range_ge_0_10 | coding_or_protein_altering_differentiated | splice_related_differentiated | regulatory_or_promoter_differentiated | overlapping_UniProt_feature_differentiated | max_continental_AF_range |
|---|---|---|---|---|---|---|---|
| ABCB1 | 5702 | 50 | 0 | 0 | 0 | 0 | 0.20742826 |
| CES2 | 1306 | 27 | 0 | 0 | 5 | 0 | 0.23437919 |
| HLA-B | 4104 | 50 | 0 | 0 | 3 | 0 | 0.31313131 |
| NAT2 | 1963 | 50 | 2 | 0 | 7 | 0 | 0.1986532 |
| NR1I2 | 2089 | 50 | 0 | 0 | 3 | 0 | 0.21717172 |
| NR1I3 | 1447 | 50 | 0 | 0 | 5 | 0 | 0.2161022 |
| SLCO1B1 | 5972 | 50 | 0 | 0 | 0 | 0 | 0.25855904 |
| UGT1A1 | 1789 | 49 | 0 | 0 | 2 | 0 | 0.24667024 |

#### Supplementary Table S1 - panel B

| gene_symbol | top_differentiated_modules | top_differentiated_consequences | annotation_interpretation |
|---|---|---|---|
| ABCB1 | gene_body;<br>intronic_regulation_or_haplotype:27;<br>downstream_25kb:21;<br>gene_body:1;<br>3_prime_UTR_regulation;<br>gene_body:1 | intron_variant:27;downstream_gene_variant:22;3_prime_UTR_variant:1 | mechanistic_hypothesis_generation_only;<br>population differentiation does not establish drug-response causality |
| CES2 | distal_upstream_2_to_25kb:10;<br>downstream_25kb:8;<br>gene_body;<br>intronic_regulation_or_haplotype:4;<br>annotated_regulatory_element;<br>distal_upstream_2_to_25kb:2;<br>proximal_promoter_2kb:1;<br>annotated_regulatory_element;<br>gene_body:1;<br>annotated_regulatory_element;<br>downstream_25kb:1 | upstream_gene_variant:14;downstream_gene_variant:9;intron_variant:4 | mechanistic_hypothesis_generation_only;<br>population differentiation does not establish drug-response causality |
| HLA-B | downstream_25kb:44;<br>annotated_regulatory_element;<br>downstream_25kb:3;<br>gene_body:2;<br>distal_upstream_2_to_25kb:1 | downstream_gene_variant:47;upstream_gene_variant:3 | mechanistic_hypothesis_generation_only;<br>population differentiation does not establish drug-response causality |
| NAT2 | downstream_25kb:25;<br>distal_upstream_2_to_25kb:10;<br>gene_body;<br>intronic_regulation_or_haplotype:6;<br>annotated_regulatory_element;<br>distal_upstream_2_to_25kb:5;<br>coding_synonymous;<br>gene_body:2;<br>annotated_regulatory_element;<br>downstream_25kb:2 | downstream_gene_variant:27;<br>upstream_gene_variant:15;<br>intron_variant:6;<br>synonymous_variant:2 | mechanistic_hypothesis_generation_only;<br>population differentiation does not establish drug-response causality |
| NR1I2 | gene_body;<br>intronic_regulation_or_haplotype:35;<br>downstream_25kb:4;<br>distal_upstream_2_to_25kb:3;<br>3_prime_UTR_regulation;<br>gene_body:3;<br>proximal_promoter_2kb:2;<br>gene_body:2;<br>annotated_regulatory_element;<br>downstream_25kb:1 | intron_variant:35;<br>upstream_gene_variant:7;<br>downstream_gene_variant:5;<br>3_prime_UTR_variant:3 | mechanistic_hypothesis_generation_only;<br>population differentiation does not establish drug-response causality |
| NR1I3 | distal_upstream_2_to_25kb:26;<br>downstream_25kb:10;<br>gene_body;<br>intronic_regulation_or_haplotype:9;<br>proximal_promoter_2kb:3;<br>annotated_regulatory_element;<br>downstream_25kb:2 | upstream_gene_variant:29;downstream_gene_variant:12;intron_variant:9 | mechanistic_hypothesis_generation_only;<br>population differentiation does not establish drug-response causality |
| SLCO1B1 | gene_body:25;<br>gene_body;<br>intronic_regulation_or_haplotype:23;<br>downstream_25kb:2 | intron_variant:23;<br>intron_variant;<br>NMD_transcript_variant:23;<br>downstream_gene_variant:3;<br>3_prime_UTR_variant;<br>NMD_transcript_variant:1 | mechanistic_hypothesis_generation_only;<br>population differentiation does not establish drug-response causality |
| UGT1A1 | downstream_25kb:26;<br>distal_upstream_2_to_25kb:15;<br>gene_body;<br>intronic_regulation_or_haplotype:4;<br>proximal_promoter_2kb:2;<br>3_prime_UTR_regulation;<br>gene_body:2 | downstream_gene_variant:26;<br>upstream_gene_variant:17;<br>intron_variant:4;<br>3_prime_UTR_variant:2 | mechanistic_hypothesis_generation_only;<br>population differentiation does not establish drug-response causality |

### Supplementary Table S2. NAT2 genotype-inferred acetylator categories by population

Phenotypes are inferred from NAT2 genotype and are not measured pharmacokinetic outcomes.

#### Supplementary Table S2 - panel A

| sample_set | population_code | predicted_acetylator_phenotype | individual_count | individual_total | frequency |
|---|---|---|---|---|---|
| continental_AFR | ESN | indeterminate | 6 | 99 | 0.06060606060606061 |
| continental_AFR | ESN | intermediate_metabolizer | 35 | 99 | 0.35353535353535354 |
| continental_AFR | ESN | poor_metabolizer | 48 | 99 | 0.48484848484848486 |
| continental_AFR | ESN | rapid_metabolizer | 10 | 99 | 0.10101010101010101 |
| continental_AFR | GWD | indeterminate | 15 | 113 | 0.13274336283185842 |
| continental_AFR | GWD | intermediate_metabolizer | 38 | 113 | 0.336283185840708 |
| continental_AFR | GWD | poor_metabolizer | 53 | 113 | 0.4690265486725664 |
| continental_AFR | GWD | rapid_metabolizer | 7 | 113 | 0.061946902654867256 |
| continental_AFR | LWK | indeterminate | 5 | 99 | 0.050505050505050504 |
| continental_AFR | LWK | intermediate_metabolizer | 35 | 99 | 0.35353535353535354 |
| continental_AFR | LWK | poor_metabolizer | 53 | 99 | 0.5353535353535354 |
| continental_AFR | LWK | rapid_metabolizer | 6 | 99 | 0.06060606060606061 |
| continental_AFR | MSL | indeterminate | 2 | 85 | 0.023529411764705882 |
| continental_AFR | MSL | intermediate_metabolizer | 44 | 85 | 0.5176470588235295 |
| continental_AFR | MSL | poor_metabolizer | 27 | 85 | 0.3176470588235294 |
| continental_AFR | MSL | rapid_metabolizer | 12 | 85 | 0.1411764705882353 |
| continental_AFR | YRI | indeterminate | 7 | 108 | 0.06481481481481481 |
| continental_AFR | YRI | intermediate_metabolizer | 57 | 108 | 0.5277777777777778 |
| continental_AFR | YRI | poor_metabolizer | 33 | 108 | 0.3055555555555556 |
| continental_AFR | YRI | rapid_metabolizer | 11 | 108 | 0.10185185185185185 |
| diaspora_context | ACB | indeterminate | 5 | 96 | 0.052083333333333336 |
| diaspora_context | ACB | intermediate_metabolizer | 43 | 96 | 0.4479166666666667 |
| diaspora_context | ACB | poor_metabolizer | 40 | 96 | 0.4166666666666667 |
| diaspora_context | ACB | rapid_metabolizer | 8 | 96 | 0.08333333333333333 |
| diaspora_context | ASW | indeterminate | 1 | 61 | 0.01639344262295082 |
| diaspora_context | ASW | intermediate_metabolizer | 22 | 61 | 0.36065573770491804 |
| diaspora_context | ASW | poor_metabolizer | 33 | 61 | 0.5409836065573771 |
| diaspora_context | ASW | rapid_metabolizer | 5 | 61 | 0.08196721311475409 |

### Supplementary Table S3. Conditional noncore NAT2 background heterogeneity across populations

The five global permutation tests use individual-carrier clustering, with a separate global BH correction family.

#### Supplementary Table S3 - panel A

| core_allele | haplotype_count | carrier_individual_count | homozygous_carrier_count | distinct_noncore_backgrounds | observed_generalized_jsd | permutation_count | extreme_permutation_count | empirical_p | bh_q_global_5 | significant_global_bh_0_05 |
|---|---|---|---|---|---|---|---|---|---|---|
| *5 | 285 | 242 | 43 | 34 | 0.465685775854 | 50000 | 0 | 1.9999600008e-05 | 9.999800004e-05 | YES |
| *4 | 233 | 203 | 30 | 44 | 0.55187523702 | 50000 | 741 | 0.0148397032059 | 0.0247328386766 | YES |
| *6 | 182 | 166 | 16 | 27 | 0.59770134022 | 50000 | 6 | 0.000139997200056 | 0.00034999300014 | YES |
| *14 | 111 | 106 | 5 | 10 | 0.249656938388 | 50000 | 9113 | 0.182276354473 | 0.182276354473 | NO |
| *1 | 80 | 78 | 2 | 17 | 0.632302324593 | 50000 | 6541 | 0.130837383252 | 0.163546729065 | NO |

#### Supplementary Table S3 - panel B

| core_allele | null_mean | null_sd | null_q95 | null_q99 | mc_se_approx | minimum_possible_empirical_p |
|---|---|---|---|---|---|---|
| *5 | 0.330931311697 | 0.02609247292 | 0.375666589753 | 0.396563195227 | 1.9999400015e-05 | 1.9999600008e-05 |
| *4 | 0.490561467002 | 0.026217067034 | 0.535636312677 | 0.55628300538 | 0.000540725328255 | 1.9999600008e-05 |
| *6 | 0.423766893085 | 0.040230954416 | 0.492391320048 | 0.5248797698 | 5.29102639086e-05 | 1.9999600008e-05 |
| *14 | 0.216328719517 | 0.0387919222944 | 0.286702543638 | 0.324905377182 | 0.00172654976238 | 1.9999600008e-05 |
| *1 | 0.56198494007 | 0.0618122905234 | 0.670595605283 | 0.724441638934 | 0.00150808944073 | 1.9999600008e-05 |

### Supplementary Table S4. NAT2 core-matching and acetylator-inference completeness

Analytic denominators, function uncertainty and phenotype indeterminacy are displayed explicitly.

#### Supplementary Table S4 - panel A

| analysis_scope | population_group | individual_count | haplotype_count | exact_core_definition_haplotypes | unmatched_core_haplotypes | exact_core_call_fraction | decreased_function_haplotypes | increased_function_haplotypes |
|---|---|---|---|---|---|---|---|---|
| primary_continental | ESN | 99 | 198 | 198 | 0 | 1 | 134 | 58 |
| primary_continental | GWD | 113 | 226 | 225 | 1 | 0.995575221239 | 154 | 57 |
| primary_continental | LWK | 99 | 198 | 198 | 0 | 1 | 145 | 48 |
| primary_continental | MSL | 85 | 170 | 170 | 0 | 1 | 99 | 69 |
| primary_continental | YRI | 108 | 216 | 216 | 0 | 1 | 128 | 81 |
| primary_continental | ALL_5_CONTINENTAL | 504 | 1008 | 1007 | 1 | 0.999007936508 | 660 | 313 |
| secondary_diaspora_context | ACB | 96 | 192 | 192 | 0 | 1 | 126 | 61 |
| secondary_diaspora_context | ASW | 61 | 122 | 122 | 0 | 1 | 88 | 32 |
| secondary_diaspora_context | ALL_DIASPORA_CONTEXT | 157 | 314 | 314 | 0 | 1 | 214 | 93 |
| all_contexts | ALL_661 | 661 | 1322 | 1321 | 1 | 0.999243570348 | 874 | 406 |

#### Supplementary Table S4 - panel B

| analysis_scope | uncertain_function_haplotypes | unknown_function_haplotypes | function_resolved_haplotypes | function_resolved_fraction | rapid_metabolizer_individuals | intermediate_metabolizer_individuals | poor_metabolizer_individuals | indeterminate_individuals |
|---|---|---|---|---|---|---|---|---|
| primary_continental | 6 | 0 | 192 | 0.969696969697 | 10 | 35 | 48 | 6 |
| primary_continental | 14 | 1 | 211 | 0.933628318584 | 7 | 38 | 53 | 15 |
| primary_continental | 5 | 0 | 193 | 0.974747474747 | 6 | 35 | 53 | 5 |
| primary_continental | 2 | 0 | 168 | 0.988235294118 | 12 | 44 | 27 | 2 |
| primary_continental | 7 | 0 | 209 | 0.967592592593 | 11 | 57 | 33 | 7 |
| primary_continental | 34 | 1 | 973 | 0.965277777778 | 46 | 209 | 214 | 35 |
| secondary_diaspora_context | 5 | 0 | 187 | 0.973958333333 | 8 | 43 | 40 | 5 |
| secondary_diaspora_context | 2 | 0 | 120 | 0.983606557377 | 5 | 22 | 33 | 1 |
| secondary_diaspora_context | 7 | 0 | 307 | 0.977707006369 | 13 | 65 | 73 | 6 |
| all_contexts | 41 | 1 | 1280 | 0.968229954614 | 59 | 274 | 287 | 41 |

#### Supplementary Table S4 - panel C

| analysis_scope | phenotype_determinate_individuals | phenotype_determinate_fraction | indeterminate_with_unmatched_core | indeterminate_without_unmatched_core |
|---|---|---|---|---|
| primary_continental | 93 | 0.939393939394 | 0 | 6 |
| primary_continental | 98 | 0.867256637168 | 1 | 14 |
| primary_continental | 94 | 0.949494949495 | 0 | 5 |
| primary_continental | 83 | 0.976470588235 | 0 | 2 |
| primary_continental | 101 | 0.935185185185 | 0 | 7 |
| primary_continental | 469 | 0.930555555556 | 1 | 34 |
| secondary_diaspora_context | 91 | 0.947916666667 | 0 | 5 |
| secondary_diaspora_context | 60 | 0.983606557377 | 0 | 1 |
| secondary_diaspora_context | 151 | 0.96178343949 | 0 | 6 |
| all_contexts | 620 | 0.937972768533 | 1 | 40 |

### Supplementary Table S5. NAT2 core-group integrated global summaries

Only core groups meeting the predefined representation criteria were eligible for the five-population global test.

#### Supplementary Table S5 - panel A

| core_allele | core_function | primary_continental_haplotype_count | distinct_35_site_backgrounds | effective_backgrounds_shannon | inverse_simpson | dominant_background_fraction | five_population_inferential_eligibility | minimum_population_haplotype_count |
|---|---|---|---|---|---|---|---|---|
| *5 | Decreased function | 285 | 34 | 11.536136660407962 | 7.13188164017912 | 0.23157894736842105 | YES | 40 |
| *4 | Increased function | 233 | 44 | 9.125765627825972 | 4.100687363093889 | 0.4206008583690987 | YES | 34 |
| *6 | Decreased function | 182 | 27 | 11.557597488820043 | 6.512780180888715 | 0.33516483516483514 | YES | 29 |
| *14 | Decreased function | 111 | 10 | 3.086919261593187 | 2.195082843399252 | 0.6306306306306306 | YES | 12 |
| *1 | Increased function | 80 | 17 | 8.708201430648765 | 6.095238095238097 | 0.2875 | YES | 12 |
| *39 | Decreased function | 30 | 4 | 1.7011489969890363 | 1.319648093841642 | 0.8666666666666667 | NO | 2 |
| *7 | Decreased function | 25 | 6 | 4.415825582444111 | 3.7425149700598803 | 0.36 | NO | 2 |
| *44 | Uncertain function | 20 | 6 | 3.782239457341191 | 2.7777777777777772 | 0.55 | NO | 0 |

#### Supplementary Table S5 - panel B

| core_allele | minimum_population_carrier_individual_count | global_generalized_jsd | global_empirical_p | global_bh_q | global_bh_significant | bh_significant_pairwise_comparisons |
|---|---|---|---|---|---|---|
| *5 | 36 | 0.465685775854 | 1.9999600008e-05 | 9.999800004e-05 | YES | 4 |
| *4 | 29 | 0.55187523702 | 0.0148397032059 | 0.0247328386766 | YES | 0 |
| *6 | 25 | 0.59770134022 | 0.000139997200056 | 0.00034999300014 | YES | 2 |
| *14 | 11 | 0.249656938388 | 0.182276354473 | 0.182276354473 | NO | 0 |
| *1 | 12 | 0.632302324593 | 0.130837383252 | 0.163546729065 | NO | 0 |
| *39 | 2 | | | | NOT_TESTED | 0 |
| *7 | 2 | | | | NOT_TESTED | 0 |
| *44 | 0 | | | | NOT_TESTED | 0 |

### Supplementary Table S6. All 50 pairwise NAT2 background-heterogeneity contrasts

All 50 pairwise comparisons belong to one separately BH-adjusted family. Source identifiers repeat in each panel.

#### Supplementary Table S6 - panel A

Fields core_allele, population_a, population_b, haplotypes_a

| core_allele | population_a | population_b | haplotypes_a | haplotypes_b | carrier_individuals_a | carrier_individuals_b | homozygous_carriers_a | homozygous_carriers_b | distinct_noncore_backgrounds_allele_union | observed_pairwise_jsd |
|---|---|---|---|---|---|---|---|---|---|---|
| *5 | ESN | GWD | 53 | 73 | 46 | 59 | 7 | 14 | 34 | 0.299930928714 |
| *5 | ESN | LWK | 53 | 68 | 46 | 58 | 7 | 10 | 34 | 0.324841066402 |
| *5 | ESN | MSL | 53 | 40 | 46 | 36 | 7 | 4 | 34 | 0.250719239452 |
| *5 | ESN | YRI | 53 | 51 | 46 | 43 | 7 | 8 | 34 | 0.227860503251 |
| *5 | GWD | LWK | 73 | 68 | 59 | 58 | 14 | 10 | 34 | 0.259474603332 |
| *5 | GWD | MSL | 73 | 40 | 59 | 36 | 14 | 4 | 34 | 0.204719594266 |
| *5 | GWD | YRI | 73 | 51 | 59 | 43 | 14 | 8 | 34 | 0.212832519367 |
| *5 | LWK | MSL | 68 | 40 | 58 | 36 | 10 | 4 | 34 | 0.186072309981 |
| *5 | LWK | YRI | 68 | 51 | 58 | 43 | 10 | 8 | 34 | 0.299028372165 |
| *5 | MSL | YRI | 40 | 51 | 36 | 43 | 4 | 8 | 34 | 0.226561781459 |
| *4 | ESN | GWD | 41 | 45 | 34 | 40 | 7 | 5 | 44 | 0.242723377556 |
| *4 | ESN | LWK | 41 | 34 | 34 | 29 | 7 | 5 | 44 | 0.293189916033 |
| *4 | ESN | MSL | 41 | 53 | 34 | 47 | 7 | 6 | 44 | 0.287209996561 |
| *4 | ESN | YRI | 41 | 60 | 34 | 53 | 7 | 7 | 44 | 0.260229515779 |
| *4 | GWD | LWK | 45 | 34 | 40 | 29 | 5 | 5 | 44 | 0.245521451324 |
| *4 | GWD | MSL | 45 | 53 | 40 | 47 | 5 | 6 | 44 | 0.234657460048 |
| *4 | GWD | YRI | 45 | 60 | 40 | 53 | 5 | 7 | 44 | 0.261969792607 |
| *4 | LWK | MSL | 34 | 53 | 29 | 47 | 5 | 6 | 44 | 0.281403502749 |
| *4 | LWK | YRI | 34 | 60 | 29 | 53 | 5 | 7 | 44 | 0.350533795415 |
| *4 | MSL | YRI | 53 | 60 | 47 | 53 | 6 | 7 | 44 | 0.244911192197 |
| *6 | ESN | GWD | 44 | 32 | 38 | 30 | 6 | 2 | 27 | 0.428766016922 |
| *6 | ESN | LWK | 44 | 45 | 38 | 43 | 6 | 2 | 27 | 0.286884111454 |
| *6 | ESN | MSL | 44 | 29 | 38 | 25 | 6 | 4 | 27 | 0.345089223542 |
| *6 | ESN | YRI | 44 | 32 | 38 | 30 | 6 | 2 | 27 | 0.295978195469 |
| *6 | GWD | LWK | 32 | 45 | 30 | 43 | 2 | 2 | 27 | 0.407174460714 |
| *6 | GWD | MSL | 32 | 29 | 30 | 25 | 2 | 4 | 27 | 0.188599159687 |
| *6 | GWD | YRI | 32 | 32 | 30 | 30 | 2 | 2 | 27 | 0.322734719251 |
| *6 | LWK | MSL | 45 | 29 | 43 | 25 | 2 | 4 | 27 | 0.356010050623 |
| *6 | LWK | YRI | 45 | 32 | 43 | 30 | 2 | 2 | 27 | 0.313747667667 |
| *6 | MSL | YRI | 29 | 32 | 25 | 30 | 4 | 2 | 27 | 0.181455687898 |
| *14 | ESN | GWD | 25 | 32 | 23 | 30 | 2 | 2 | 10 | 0.165840836334 |
| *14 | ESN | LWK | 25 | 17 | 23 | 17 | 2 | 0 | 10 | 0.161416748929 |
| *14 | ESN | MSL | 25 | 12 | 23 | 11 | 2 | 1 | 10 | 0.157346158199 |
| *14 | ESN | YRI | 25 | 25 | 23 | 25 | 2 | 0 | 10 | 0.0917779032506 |

| core_allele | population_a | population_b | haplotypes_a | haplotypes_b | carrier_individuals_a | carrier_individuals_b | homozygous_carriers_a | homozygous_carriers_b | distinct_noncore_backgrounds_allele_union | observed_pairwise_jsd |
|---|---|---|---|---|---|---|---|---|---|---|
| *14 | GWD | LWK | 32 | 17 | 30 | 17 | 2 | 0 | 10 | 0.181409346738 |
| *14 | GWD | MSL | 32 | 12 | 30 | 11 | 2 | 1 | 10 | 0.060968200141 |
| *14 | GWD | YRI | 32 | 25 | 30 | 25 | 2 | 0 | 10 | 0.0663363947263 |
| *14 | LWK | MSL | 17 | 12 | 17 | 11 | 0 | 1 | 10 | 0.225123324028 |
| *14 | LWK | YRI | 17 | 25 | 17 | 25 | 0 | 0 | 10 | 0.0775303090828 |
| *14 | MSL | YRI | 12 | 25 | 11 | 25 | 1 | 0 | 10 | 0.0905679462222 |
| *1 | ESN | GWD | 17 | 12 | 17 | 12 | 0 | 0 | 17 | 0.416610679735 |
| *1 | ESN | LWK | 17 | 14 | 17 | 14 | 0 | 0 | 17 | 0.361437210384 |
| *1 | ESN | MSL | 17 | 16 | 17 | 15 | 0 | 1 | 17 | 0.268212482195 |
| *1 | ESN | YRI | 17 | 21 | 17 | 20 | 0 | 1 | 17 | 0.214816668527 |
| *1 | GWD | LWK | 12 | 14 | 12 | 14 | 0 | 0 | 17 | 0.423907067483 |
| *1 | GWD | MSL | 12 | 16 | 12 | 15 | 0 | 1 | 17 | 0.336709557529 |
| *1 | GWD | YRI | 12 | 21 | 12 | 20 | 0 | 1 | 17 | 0.244725277544 |
| *1 | LWK | MSL | 14 | 16 | 14 | 15 | 0 | 1 | 17 | 0.455780125484 |
| *1 | LWK | YRI | 14 | 21 | 14 | 20 | 0 | 1 | 17 | 0.269952860033 |
| *1 | MSL | YRI | 16 | 21 | 15 | 20 | 1 | 1 | 17 | 0.272226458695 |

#### Supplementary Table S6 - panel B

Fields core_allele, population_a, population_b, permutation_count

| core_allele | population_a | population_b | permutation_count | extreme_permutation_count | empirical_p | bh_q_pairwise_50 | significant_pairwise_bh_0_05 | null_mean | null_sd |
|---|---|---|---|---|---|---|---|---|---|
| *5 | ESN | GWD | 50000 | 1 | 3.9999200016e-05 | 0.0019999600008 | YES | 0.132052498507 | 0.0320739431847 |
| *5 | ESN | LWK | 50000 | 8 | 0.000179996400072 | 0.0019999600008 | YES | 0.16361327759 | 0.035029543594 |
| *5 | ESN | MSL | 50000 | 1132 | 0.0226595468091 | 0.0905366508054 | NO | 0.162572706246 | 0.0400340137294 |
| *5 | ESN | YRI | 50000 | 10016 | 0.20033599328 | 0.302052782474 | NO | 0.200462487032 | 0.0342179638093 |
| *5 | GWD | LWK | 50000 | 8 | 0.000179996400072 | 0.0019999600008 | YES | 0.122883242657 | 0.030298205198 |
| *5 | GWD | MSL | 50000 | 999 | 0.019999600008 | 0.0905366508054 | NO | 0.126106532052 | 0.0338947429537 |
| *5 | GWD | YRI | 50000 | 1849 | 0.0369992600148 | 0.114719927824 | NO | 0.152726272803 | 0.0305549645125 |
| *5 | LWK | MSL | 50000 | 7850 | 0.157016859663 | 0.290771962339 | NO | 0.151669790889 | 0.0342902148327 |
| *5 | LWK | YRI | 50000 | 104 | 0.00209995800084 | 0.017499650007 | YES | 0.185084138341 | 0.0337440706898 |
| *5 | MSL | YRI | 50000 | 10011 | 0.20023599528 | 0.302052782474 | NO | 0.200161781544 | 0.0334709234882 |
| *4 | ESN | GWD | 50000 | 4410 | 0.0882182356353 | 0.200474251385 | NO | 0.20456329208 | 0.0260594759168 |
| *4 | ESN | LWK | 50000 | 2064 | 0.0412991740165 | 0.114719927824 | NO | 0.235963508312 | 0.0284101192952 |
| *4 | ESN | MSL | 50000 | 611 | 0.0122397552049 | 0.0679986400272 | NO | 0.208285856119 | 0.0297900793228 |
| *4 | ESN | YRI | 50000 | 10269 | 0.205395892082 | 0.302052782474 | NO | 0.236026846228 | 0.0305174125693 |
| *4 | GWD | LWK | 50000 | 18976 | 0.379532409352 | 0.485790284194 | NO | 0.238098865769 | 0.0275201848048 |
| *4 | GWD | MSL | 50000 | 4610 | 0.0922181556369 | 0.200474251385 | NO | 0.188896219249 | 0.0331513605633 |
| *4 | GWD | YRI | 50000 | 9694 | 0.193896122078 | 0.302052782474 | NO | 0.235254081071 | 0.0312319293636 |
| *4 | LWK | MSL | 50000 | 3308 | 0.0661786764265 | 0.174154411649 | NO | 0.227360973856 | 0.033477843621 |
| *4 | LWK | YRI | 50000 | 1815 | 0.0363192736145 | 0.114719927824 | NO | 0.289119069718 | 0.0322951374976 |
| *4 | MSL | YRI | 50000 | 10227 | 0.204555908882 | 0.302052782474 | NO | 0.223213479765 | 0.0280427405623 |
| *6 | ESN | GWD | 50000 | 4 | 9.999800004e-05 | 0.0019999600008 | YES | 0.197135756319 | 0.0466908614515 |
| *6 | ESN | LWK | 50000 | 559 | 0.0111997760045 | 0.0679986400272 | NO | 0.173723027427 | 0.0408374588379 |
| *6 | ESN | MSL | 50000 | 1176 | 0.0235395292094 | 0.0905366508054 | NO | 0.229576516905 | 0.0503584296896 |
| *6 | ESN | YRI | 50000 | 5186 | 0.103737925241 | 0.216120677586 | NO | 0.236915936314 | 0.0448432121472 |
| *6 | GWD | LWK | 50000 | 9 | 0.00019999600008 | 0.0019999600008 | YES | 0.186558834217 | 0.0461684477805 |
| *6 | GWD | MSL | 50000 | 22889 | 0.457790844183 | 0.508656493537 | NO | 0.191061802223 | 0.0441326566795 |
| *6 | GWD | YRI | 50000 | 1701 | 0.0340393192136 | 0.114719927824 | NO | 0.224820112789 | 0.0479817676225 |
| *6 | LWK | MSL | 50000 | 529 | 0.0105997880042 | 0.0679986400272 | NO | 0.217339281656 | 0.050578677021 |
| *6 | LWK | YRI | 50000 | 1973 | 0.0394792104158 | 0.114719927824 | NO | 0.224872911088 | 0.0452797414645 |
| *6 | MSL | YRI | 50000 | 39321 | 0.786424271515 | 0.786424271515 | NO | 0.228751706797 | 0.0582272259275 |
| *14 | ESN | GWD | 50000 | 3983 | 0.0796784064319 | 0.19919601608 | NO | 0.117148655788 | 0.0289399198518 |
| *14 | ESN | LWK | 50000 | 33116 | 0.662326753465 | 0.688435210888 | NO | 0.187192101938 | 0.0426053504413 |
| *14 | ESN | MSL | 50000 | 22854 | 0.457090858183 | 0.508656493537 | NO | 0.167024457679 | 0.060182096644 |
| *14 | ESN | YRI | 50000 | 33733 | 0.67466650667 | 0.688435210888 | NO | 0.11273237964 | 0.0327535143114 |
| *14 | GWD | LWK | 50000 | 971 | 0.0194396112078 | 0.0905366508054 | NO | 0.100538243745 | 0.0325956598697 |
| *14 | GWD | MSL | 50000 | 31736 | 0.634727305454 | 0.675241814313 | NO | 0.0747281257802 | 0.0452389772814 |
| *14 | GWD | YRI | 50000 | 14839 | 0.296794064119 | 0.412213977943 | NO | 0.047345329485 | 0.028285876518 |
| *14 | LWK | MSL | 50000 | 6593 | 0.131877362453 | 0.253610312409 | NO | 0.14460226273 | 0.0698102345186 |

| core_allele | population_a | population_b | permutation_count | extreme_permutation_count | empirical_p | bh_q_pairwise_50 | significant_pairwise_bh_0_05 | null_mean | null_sd |
|---|---|---|---|---|---|---|---|---|---|
| *14 | LWK | YRI | 50000 | 25426 | 0.508529829403 | 0.552749814569 | NO | 0.0931238923161 | 0.0425484148205 |
| *14 | MSL | YRI | 50000 | 13203 | 0.264074718506 | 0.377249597865 | NO | 0.0609517674543 | 0.0578344593806 |
| *1 | ESN | GWD | 50000 | 4289 | 0.0857982840343 | 0.200474251385 | NO | 0.273581796645 | 0.0989361803377 |
| *1 | ESN | LWK | 50000 | 8829 | 0.176596468071 | 0.302052782474 | NO | 0.281394912043 | 0.0887587929247 |
| *1 | ESN | MSL | 50000 | 21438 | 0.428771424572 | 0.49857142392 | NO | 0.26034559504 | 0.0868316690318 |
| *1 | ESN | YRI | 50000 | 16696 | 0.333933321334 | 0.449254172811 | NO | 0.19618345151 | 0.0695528232295 |
| *1 | GWD | LWK | 50000 | 10184 | 0.203695926081 | 0.302052782474 | NO | 0.347560987331 | 0.0859916411571 |
| *1 | GWD | MSL | 50000 | 19431 | 0.388632227355 | 0.485790284194 | NO | 0.316224666464 | 0.0834867024594 |
| *1 | GWD | YRI | 50000 | 21028 | 0.420571588568 | 0.49857142392 | NO | 0.241085189424 | 0.0715047020455 |
| *1 | LWK | MSL | 50000 | 5640 | 0.112817743645 | 0.22563548729 | NO | 0.360574569172 | 0.0794099830833 |
| *1 | LWK | YRI | 50000 | 17071 | 0.341433171337 | 0.449254172811 | NO | 0.250657058751 | 0.0713631940508 |
| *1 | MSL | YRI | 50000 | 20770 | 0.415411691766 | 0.49857142392 | NO | 0.26338794065 | 0.0659985157345 |

#### Supplementary Table S6 - panel C

Fields core_allele, population_a, population_b, null_q95

| core_allele | population_a | population_b | null_q95 | null_q99 | mc_se_approx | minimum_possible_empirical_p |
|---|---|---|---|---|---|---|
| *5 | ESN | GWD | 0.189183113438 | 0.218515514193 | 2.82831399049e-05 | 1.9999600008e-05 |
| *5 | ESN | LWK | 0.224967883197 | 0.25428963304 | 5.9993399997e-05 | 1.9999600008e-05 |
| *5 | ESN | MSL | 0.232747915462 | 0.269221288727 | 0.000665517074679 | 1.9999600008e-05 |
| *5 | ESN | YRI | 0.262255583091 | 0.295516437871 | 0.00178996245274 | 1.9999600008e-05 |
| *5 | GWD | LWK | 0.176007443815 | 0.200632377195 | 5.9993399997e-05 | 1.9999600008e-05 |
| *5 | GWD | MSL | 0.186389837447 | 0.217797688813 | 0.00062608663974 | 1.9999600008e-05 |
| *5 | GWD | YRI | 0.206769917109 | 0.236483220462 | 0.000844151670982 | 1.9999600008e-05 |
| *5 | LWK | MSL | 0.212182812999 | 0.24275323294 | 0.00162702131668 | 1.9999600008e-05 |
| *5 | LWK | YRI | 0.244151162764 | 0.273710810618 | 0.000204719626177 | 1.9999600008e-05 |
| *5 | MSL | YRI | 0.261223464558 | 0.294759110555 | 0.0017896275518 | 1.9999600008e-05 |
| *4 | ESN | GWD | 0.25256319926 | 0.281394771385 | 0.0012683388337 | 1.9999600008e-05 |
| *4 | ESN | LWK | 0.289401702557 | 0.313622917767 | 0.00088986246563 | 1.9999600008e-05 |
| *4 | ESN | MSL | 0.262250953897 | 0.290310563072 | 0.000491725569874 | 1.9999600008e-05 |
| *4 | ESN | YRI | 0.289129949027 | 0.315742914646 | 0.00180668290242 | 1.9999600008e-05 |
| *4 | GWD | LWK | 0.285768579667 | 0.310119300425 | 0.00217017441671 | 1.9999600008e-05 |
| *4 | GWD | MSL | 0.24681698222 | 0.273758265445 | 0.0012939265293 | 1.9999600008e-05 |
| *4 | GWD | YRI | 0.289429903741 | 0.313864143783 | 0.00176803444522 | 1.9999600008e-05 |
| *4 | LWK | MSL | 0.28747883886 | 0.316489740876 | 0.00111173578926 | 1.9999600008e-05 |
| *4 | LWK | YRI | 0.344802918524 | 0.371465547307 | 0.000836653859 | 1.9999600008e-05 |
| *4 | MSL | YRI | 0.273709250142 | 0.301176520433 | 0.00180393755342 | 1.9999600008e-05 |
| *6 | ESN | GWD | 0.281328395759 | 0.324997082762 | 4.47182291063e-05 | 1.9999600008e-05 |
| *6 | ESN | LWK | 0.248306815279 | 0.290026384164 | 0.000470619156846 | 1.9999600008e-05 |
| *6 | ESN | MSL | 0.320734675676 | 0.368945678484 | 0.00067801121045 | 1.9999600008e-05 |
| *6 | ESN | YRI | 0.318840416709 | 0.363219157392 | 0.00136363124501 | 1.9999600008e-05 |
| *6 | GWD | LWK | 0.269586881567 | 0.31434688484 | 6.3237963699e-05 | 1.9999600008e-05 |
| *6 | GWD | MSL | 0.272632716471 | 0.31907395492 | 0.00222806383614 | 1.9999600008e-05 |
| *6 | GWD | YRI | 0.312308211201 | 0.356791451491 | 0.000810925229122 | 1.9999600008e-05 |
| *6 | LWK | MSL | 0.308422020694 | 0.357648743521 | 0.000457978662255 | 1.9999600008e-05 |
| *6 | LWK | YRI | 0.307380731417 | 0.351750002269 | 0.000870859850537 | 1.9999600008e-05 |
| *6 | MSL | YRI | 0.328581871158 | 0.383625740898 | 0.00183279992106 | 1.9999600008e-05 |
| *14 | ESN | GWD | 0.174813690722 | 0.208968479645 | 0.00121101850865 | 1.9999600008e-05 |
| *14 | ESN | LWK | 0.262472288516 | 0.328921590181 | 0.00211492577741 | 1.9999600008e-05 |
| *14 | ESN | MSL | 0.294534435862 | 0.34397485496 | 0.00222779641113 | 1.9999600008e-05 |
| *14 | ESN | YRI | 0.184458380224 | 0.237988448097 | 0.00209517169561 | 1.9999600008e-05 |
| *14 | GWD | LWK | 0.16193955109 | 0.201248818926 | 0.000617435526954 | 1.9999600008e-05 |
| *14 | GWD | MSL | 0.163922359045 | 0.220958327244 | 0.00215334120049 | 1.9999600008e-05 |
| *14 | GWD | YRI | 0.0982149292372 | 0.134441535518 | 0.0020430524886 | 1.9999600008e-05 |
| *14 | LWK | MSL | 0.275716950956 | 0.381948577618 | 0.00151316512025 | 1.9999600008e-05 |

| core_allele | population_a | population_b | null_q95 | null_q99 | mc_se_approx | minimum_possible_empirical_p |
|---|---|---|---|---|---|---|
| *14 | LWK | YRI | 0.166875717406 | 0.231424435048 | 0.00223572021311 | 1.9999600008e-05 |
| *14 | MSL | YRI | 0.184648213317 | 0.248078097022 | 0.00197147343296 | 1.9999600008e-05 |
| *1 | ESN | GWD | 0.450504473143 | 0.543797682659 | 0.00125248049712 | 1.9999600008e-05 |
| *1 | ESN | LWK | 0.448225815945 | 0.528120520829 | 0.00170532839882 | 1.9999600008e-05 |
| *1 | ESN | MSL | 0.416056942408 | 0.498732909142 | 0.00221324012078 | 1.9999600008e-05 |
| *1 | ESN | YRI | 0.324173851536 | 0.403915498036 | 0.00210911075996 | 1.9999600008e-05 |
| *1 | GWD | LWK | 0.503550556503 | 0.596342212953 | 0.00180111438707 | 1.9999600008e-05 |
| *1 | GWD | MSL | 0.468334273573 | 0.578803506267 | 0.00217987369986 | 1.9999600008e-05 |
| *1 | GWD | YRI | 0.372984171974 | 0.462664056634 | 0.00220765148396 | 1.9999600008e-05 |
| *1 | LWK | MSL | 0.507351928898 | 0.599946532241 | 0.00141483496286 | 1.9999600008e-05 |
| *1 | LWK | YRI | 0.379228578867 | 0.457229732857 | 0.00212062285099 | 1.9999600008e-05 |
| *1 | MSL | YRI | 0.378073714113 | 0.455858112334 | 0.0022038146987 | 1.9999600008e-05 |
